# Genome-wide Copy Number Variations in a Large Cohort of Bantu African Children

**DOI:** 10.1101/2020.12.24.424207

**Authors:** Feyza Yilmaz, Megan Null, David Astling, Hung-Chun Yu, Joanne Cole, Stephanie A. Santorico, Benedikt Hallgrimsson, Mange Manyama, Richard A. Spritz, Audrey E. Hendricks, Tamim H. Shaikh

## Abstract

**Background:** Copy number variations (CNVs) account for a substantial proportion of inter-individual genomic variation. However, a majority of genomic variation studies have focused on single-nucleotide variations (SNVs), with limited genome-wide analysis of CNVs in large cohorts, especially in populations that are under-represented in genetic studies including people of African descent.

**Results:** In this study, we carried out a genome-wide analysis in > 3400 healthy Bantu Africans from Tanzania using high density (> 2.5 million probes) genotyping arrays. We identified over 400000 CNVs larger than 1 kilobase (kb), for an average of 120 CNVs (SE = 2.57) per individual. We detected 866 large CNVs (≥ 300 kb), some of which overlapped genomic regions previously associated with multiple congenital anomaly syndromes, including Prader-Willi/Angelman syndrome (Type1) and 22q11.2 deletion syndrome. Furthermore, several of the common CNVs seen in our cohort (≥ 5%) overlap genes previously associated with developmental disorders.

**Conclusion:** These findings may help refine the phenotypic outcomes and penetrance of variations affecting genes and genomic regions previously implicated in diseases. Our study provides one of the largest datasets of CNVs from individuals of African ancestry, enabling improved clinical evaluation and disease association of CNVs observed in research and clinical studies in African populations.

## Background

Copy number variations (CNVs) are a class of structural variation resulting from loss or gain of genomic fragments ≥ 1 kilobase (kb). CNVs can arise from genomic rearrangements such as deletions, duplications, insertions, inversions, or translocations (1–3) and have been implicated in the etiology of Mendelian disorders as well as complex traits (4). Several pediatric disorders resulting from CNVs such as microdeletions and microduplications are characterized by the occurrence of multiple congenital anomalies, including intellectual and developmental disabilities, congenital heart defects, craniofacial dysmorphisms, or abnormalities in the development of other tissues and organs (5–7). The 22q11 deletion syndrome (previously referred to as DiGeorge/Velocardiofacial Syndrome), the Williams-Beuren syndrome resulting from a microdeletion in 7q11.23, and the 15q13.3 microdeletion syndrome are examples of disorders that belong to this group of CNV-mediated disorders (8–10). CNVs may also play a role in the etiology of common, complex diseases and traits including diabetes, asthma, HIV susceptibility, cancer, and phenotypes in immune and environmental responses (11–15).

In addition to their role in disease, CNVs account for a high level of variation between apparently healthy individuals, both within and between populations (1– 3,16,17). Nevertheless, CNVs remain largely understudied compared to single-nucleotide variations (SNVs) and are not commonly genotyped in a microarray-based analysis of genome-wide variation and association to disease phenotypes (18). In 2015, Zarrei and colleagues compiled a CNV map of the human genome and estimated that 4.8-9.5% of the human genome contributes to CNV (19). Furthermore, they identified approximately 100 genes whose loss is not associated with any severe consequences (19). However, the vast majority of CNV data derive from individuals of European descent residing in Western countries, which might cause incorrect clinical interpretation of genomic variants (20). Recently, resources such as the Genome Aggregation Database (gnomAD) have reported structural variations, including CNVs, in large cohorts of individuals of both European and non-European ancestries (21). Regardless, knowledge of the genomic landscape of CNVs remains incomplete, especially in understudied populations such as Africans.

Based on the significant role of CNVs in health and disease, it is critical to have a set of reference CNVs observed in individuals from diverse populations. These population-specific reference datasets will greatly improve clinical interpretation and can help to refine a genomic region associated with diseases (22). A recent study by Kessler and colleagues (20) demonstrated how lack of African ancestry individuals in variant databases may have resulted in mischaracterization of variants in the ClinVar and Human Gene Mutation Databases.

In this study, we have detected CNVs in > 3400 apparently healthy Bantu African children from Tanzania, using data from high-density (> 2.5 million probes) genotyping microarrays. We present a high-resolution map of CNVs ranging in size from 1 kb-3 mb (million bases), providing a useful resource of CNV genetic variation for individuals of African ancestry. Additionally, we observe large CNVs in genomic regions previously implicated in syndromes and developmental disorders.

## Results

### CNV Detection and Analysis

We identified 448337 CNVs after CNV analysis and filtering from 3463 Bantu children with no known birth defects (described in Materials and Methods, Fig. 1). After merging adjacent CNVs of the same type within a given individual (see Materials and Methods), we found a total of 416877 CNVs across all autosomes, including 355027 losses and 61850 gains (Table 1, Additional Table S1). Of these, 72205 (17.3%) CNVs were concordantly called by all three CNV calling algorithms used. The average number of CNVs per subject was 120 (min = 27, max = 1569, mean = 120.38, stdev = 151.04, IQR = 45) with a median length of 7558 nucleotides (nt) and an average length of 18145 nt (min = 1,001 nt, max = 2929312 nt). We further categorized CNVs based on their genomic size, resulting in 247314 (59.3%) CNVs 1-10 kb, 158190 (38.0%) CNVs 10-100 kb, 10282 (2.5%) CNVs 100-300 kb and 1091 (0.26%) CNVs ≥ 300 kb (Table 1). Our CNV calls were significantly enriched for the Database of Genomic Variants (DGV) Gold Standard (GS) variants compared to randomly selected CNV regions (permuted p-value < 0.001), indicating that CNV calls detected in this study are likely true positives.

**Table 1.** Number and size distribution of CNVs in Bantu Africans

| CNV Length | Number of Probes | CNV |  |  |
| --- | --- | --- | --- | --- |
|  | Count | Loss | Gain | Total |
| 1-10 ≥ kb | 5-10 | 129752 | 4878 | 134630 |
|  | 11-25 | 80315 | 14633 | 94948 |
|  | 26-50 | 11733 | 4049 | 15782 |
|  | 51-100 | 985 | 910 | 1895 |
|  | >100 | 4 | 55 | 59 |
|  | **Total** | **222789** | **24525** | **247314** |
| > 10-100 ≥ kb | 5-10 | 15900 | 1680 | 17580 |
|  | 11-25 | 45287 | 9715 | 55002 |
|  | 26-50 | 41636 | 12560 | 54196 |
|  | 51-100 | 18785 | 6590 | 25375 |
|  | >100 | 3714 | 2323 | 6037 |
|  | **Total** | **125322** | **32868** | **158190** |
| > 100-300 ≥ kb | 5-10 | 37 | 10 | 47 |
|  | 11-25 | 650 | 76 | 726 |
|  | 26-50 | 1122 | 492 | 1614 |
|  | 51-100 | 1373 | 1119 | 2492 |
|  | >100 | 3413 | 1990 | 5403 |
|  | **Total** | **6595** | **3687** | **10282** |
| ≥ 300 kb | 5-10 | 2 | 4 | 6 |
|  | 11-25 | 1 | 1 | 2 |
|  | 26-50 | 12 | 28 | 40 |
|  | 51-100 | 8 | 9 | 17 |
|  | >100 | 298 | 728 | 1026 |
|  | **Total** | **321** | **770** | **1091** |

**Figure 1:**
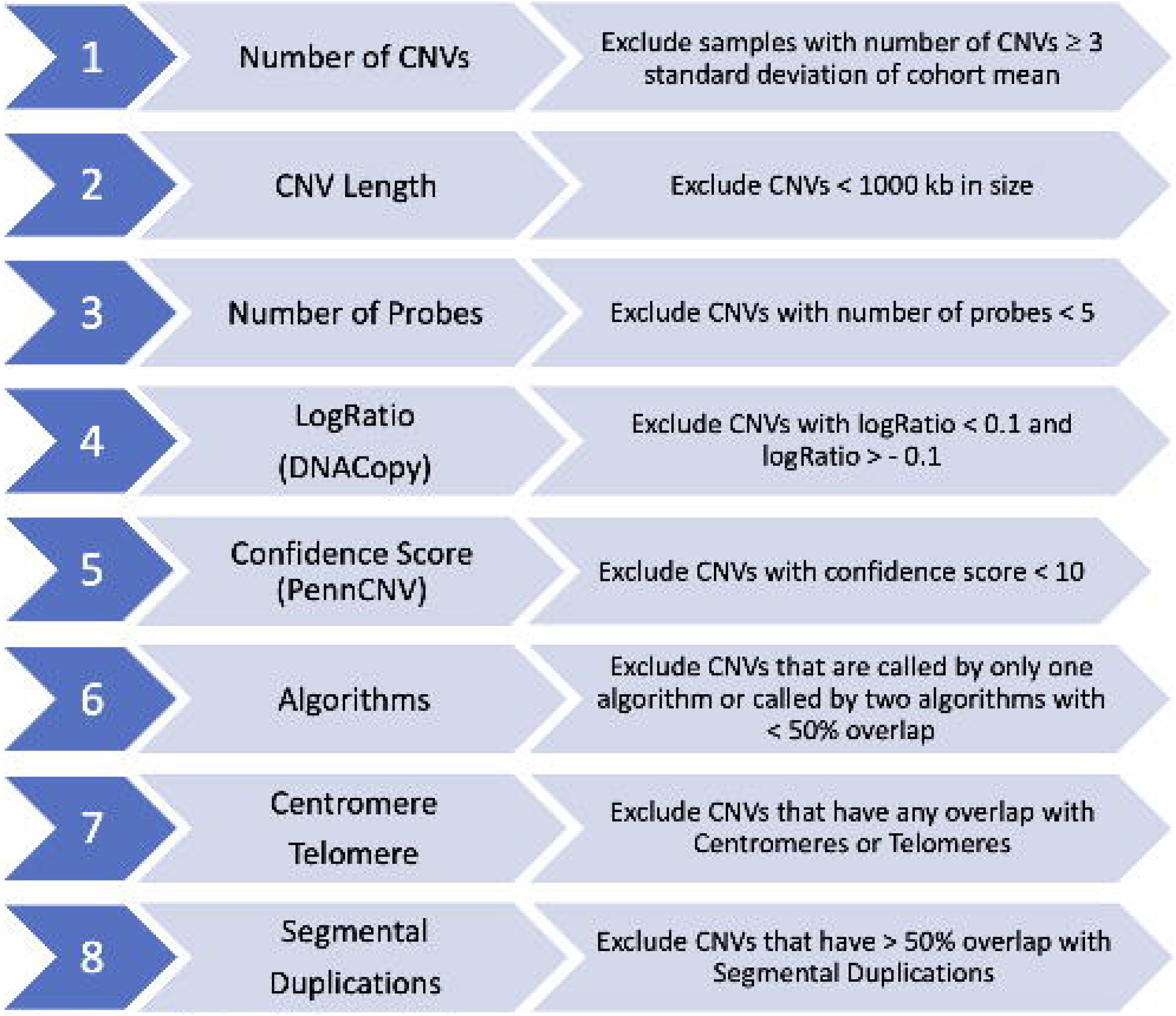
CNV Analysis and Filtering Pipeline Workflow showing the various filtering steps applied to detected CNVs in order to obtain a set of high confidence CNVs used for further analysis.

We next assembled copy number variation regions (CNVRs) by merging overlapping CNVs of the same type (loss or gain) detected in multiple individuals in the Bantu cohort (Additional Table S2). These CNVRs were further divided into 13738 loss only, 1100 gain only and 2656 with both gain and loss, for a total of 17494 CNVRs (Additional Table S2). The assembly into CNVRs further allowed us to determine that CNVs observed in our cohort covered a total of approximately 600 million nucleotides, about 20% of the genome. The distribution of CNVRs across the genome suggested that the number of CNVRs was not proportional to the size of the chromosome (Fig. 2), consistent with previous reports (19).

**Figure 2:**
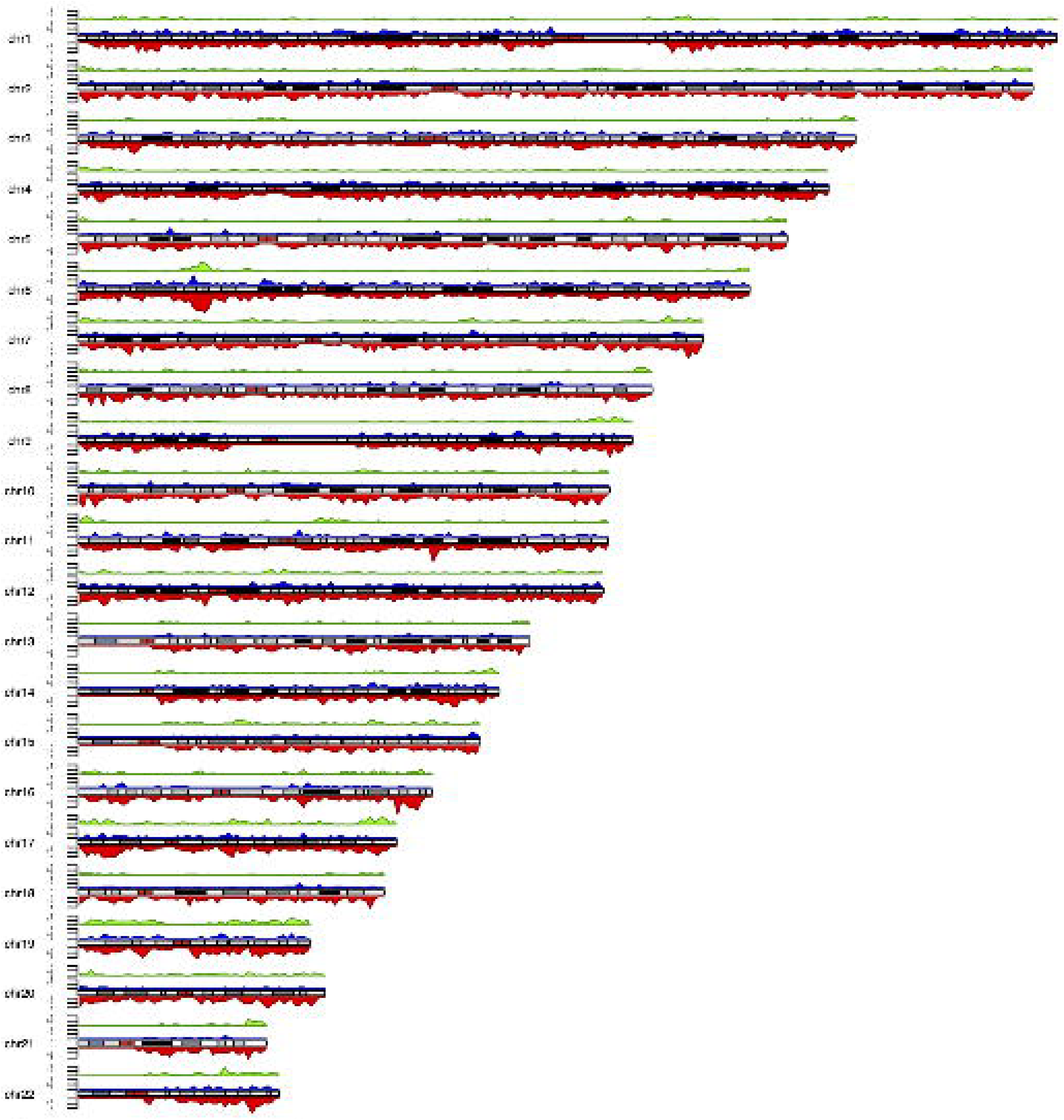
Genomic Map of CNVRs NVRs detected in our cohort are shown as colored density plots across individual chromosomes represented by ideograms. The genome was divided into 1 million equal sized windows and the number of CNVRs within each window were counted and plotted on the density plot. Color key-red: loss CNVRs, blue: gain CNVRs, green: loss and gain CNVRs. Density was calculated by dividing the genome in equal sized windows (n = 1000000) and counting the number of CNVRs overlapping each of the windows.

### Comparison to other CNV Datasets

To determine overlap with existing CNV datasets, we compared the CNVRs observed in our cohort with existing CNV databases including DGV (40418 CNVRs) (23), gnomAD (54851 CNVRs) (21,24), and current studies that focus on CNVs in different African populations (7608 CNVRs) (25) and low-mappability regions (12242 CNVRs) (26). This comparison identified 1952 (11.16%) CNVRs in our cohort overlapping all four and 10046 (57.46%) overlapping any three datasets, while a majority overlapped CNVRs in only one, two, or three of the databases (Table 2).

**Table 2.** Bantu CNVRs overlap with CNV datasets

| **CNV Datasets** | **Total CNVRs** |
| --- | --- |
| All Four | 1952 |
| Any Three | 10046 |
| Any Two | 4712 |
| DGV only | 338 |
| gnomAD only | 1 |
| Low mappability regions only | 396 |
| African CNVR | 1 |
| None | 48 |
| **Total** | **17494** |

DGV: CNVRs generated from the Database of Genomic Variants CNVs, gnomAD: CNVRs generated from Genome Aggregation Database CNVs, African CNVR: CNVRs identified by Nyangiri and colleagues (26), All Four: CNVRs observed in all four datasets. Any Three and Any Two: CNVRs from any three or two of the above datasets respectively.

Additionally, we observed 48 CNVRs in our cohort that did not overlap with any CNV datasets mentioned above (Fig. 3, Additional Table S3). These 48 CNVRs encompass a total of 209951 nt with three (very rare frequency CNVRs) overlapping genes reported to be associated with developmental disorders in the Developmental Disorders Genotype-Phenotype Database (DDG2P) (Additional Table S4). These may represent CNVRs that are either specific to the Bantu African population or that may be very rare in populations currently represented in existing CNV datasets.

**Figure 3:**
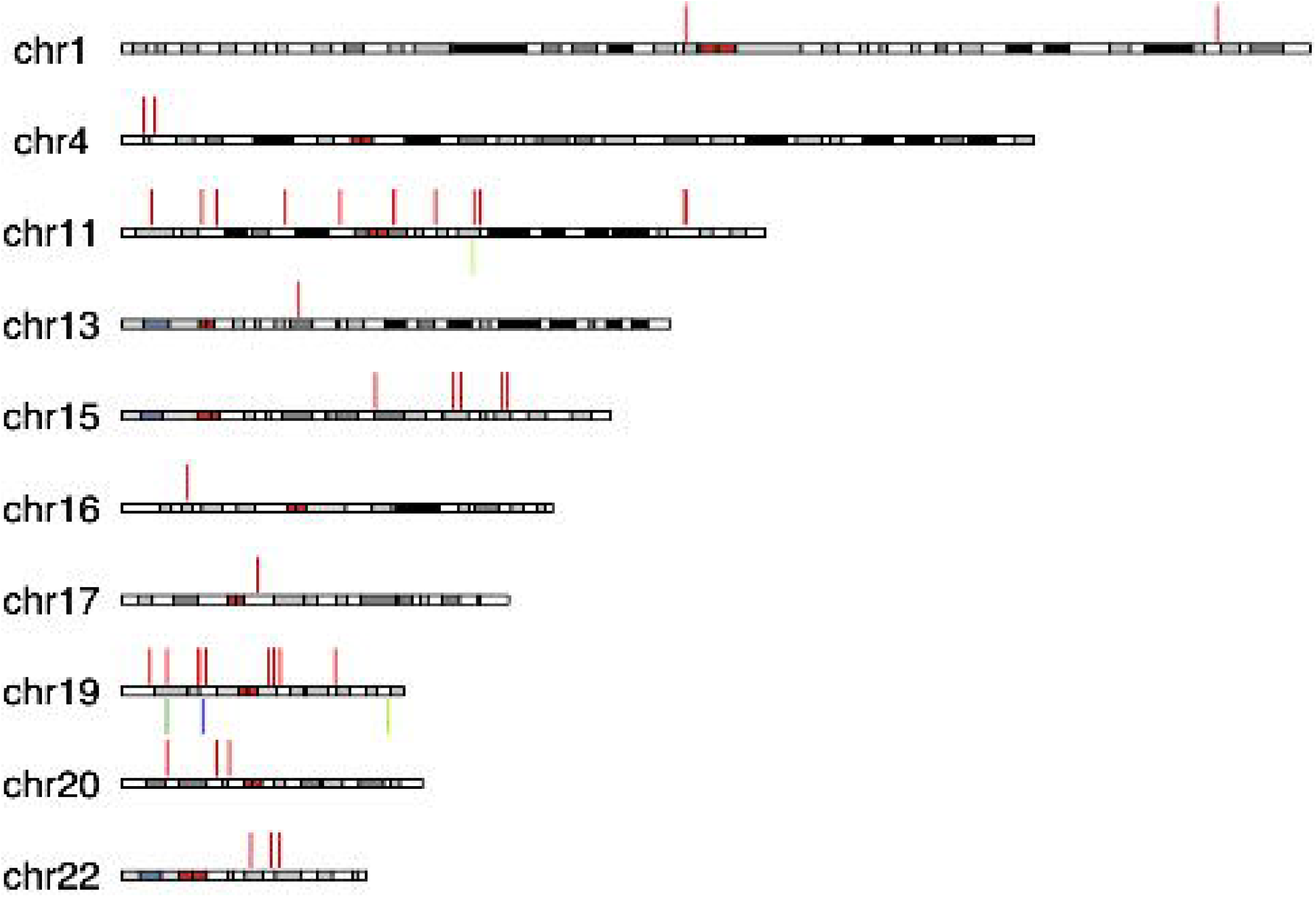
Novel Bantu CNVRs. The chromosomal locations of CNVRs detected in the Bantu cohort, which did not overlap with known CNV datasets included in our comparison analysis. Vertical, colored lines represent individual CNVRs. Color key-red: loss blue: gain, green: loss and gain

### CNVs in Regions Associated with Disease

We next wanted to determine whether CNVs observed in the Bantu cohort overlapped genes and genomic regions previously associated with disease phenotypes. Using CNVs from 2696 unrelated subjects in our cohort, we identified 121334 CNV blocks from 323667 CNV calls (Fig. 4, see Materials and Methods, Additional Table S5). We further classified CNV blocks into four categories based on how often they were observed in these 2696 unrelated individuals: a) 6913 CNV blocks observed in ≥ 5% of unrelated subjects were categorized as common; b) 24908 CNV blocks observed in 1-5% were categorized as low frequency; c) 44910 CNV blocks observed in 0.1-1% were categorized as rare; and d) 44603 CNV blocks were observed in ≤ 0.1% and were categorized as very rare; most of the very rare CNV blocks were singletons.

**Figure 4:**
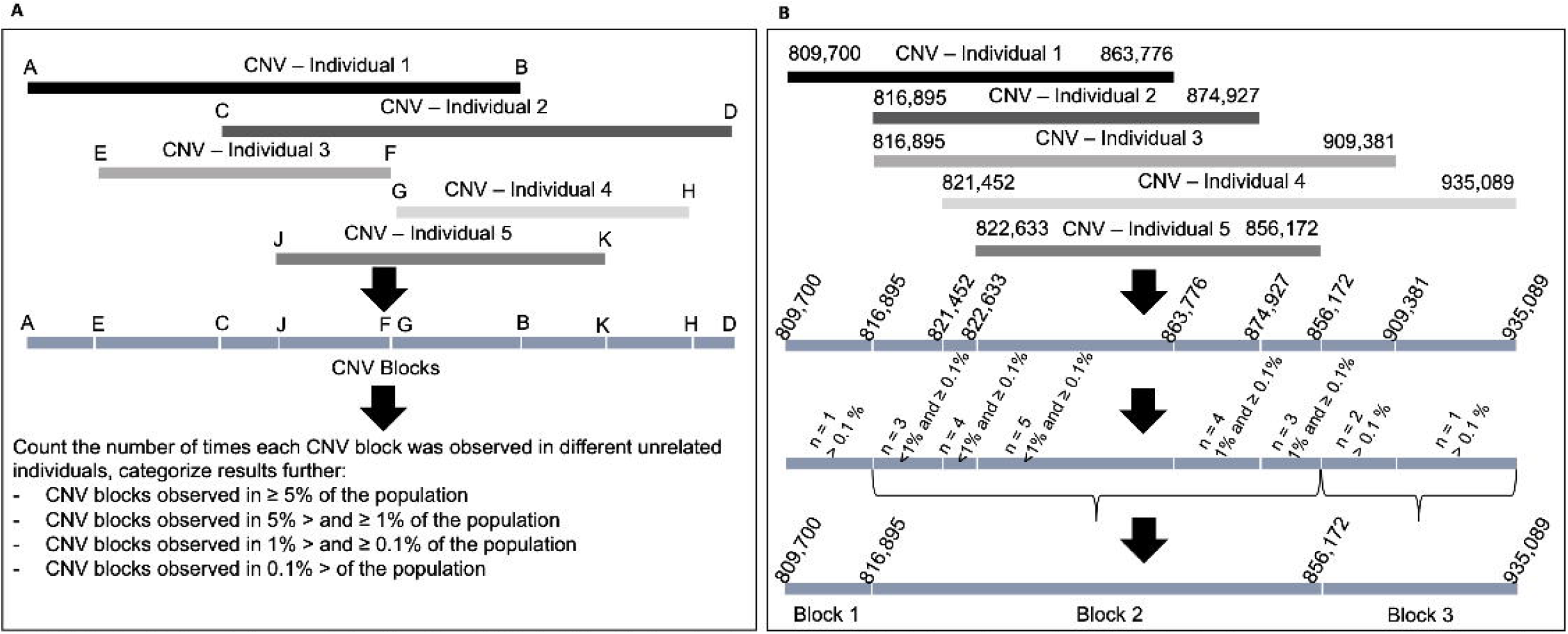
CNV blocks **a**-A schematic demonstrating the delineation of CNV blocks followed by determination of total count within the Bantu cohort and categorization based on frequency. Black and gray rectangles, A-B, C-D, E-F, G-H and J-K represent five overlapping CNVs observed in different individuals (1-5). A-K represent the start and end coordinates of the CNVs. Blue rectangles represent CNV blocks. **b**–represents an actual example of CNV block delineation from our CNV dataset.

We then determined the overlap between common (≥ 5%), low frequency (1-5%), and rare (0.1-1%) CNV blocks and genes reported to be associated with developmental disorders in the DDG2P Database (Additional Table S4). We identified 11835 CNV blocks that overlapped 1627 DDG2P genes (Additional Table S6). We used reciprocal approach to identify ≥ 50% overlap between DDG2P genes and CNV blocks, which identified 125 CNV blocks (83 loss, 21 gain, 21 loss and gain) which overlapped with 125 DDG2P genes with reciprocal overlap percentage of ≥ 50%.

Additionally, we identified 866 relatively large CNVs (≥ 300 kb) (Additional Table S7) in unrelated individuals within our cohort. We investigated whether any of these large CNVs overlap (≥ 1 bp) CNVs previously implicated in syndromes or genomic disorders catalogued in DECIPHER (DatabasE of genomiC variation and Phenotype in Humans using Ensembl Resources; Additional Table S8) (27). We identified 83 large CNVs, including 62 gain CNVs ranging in size from ~300-2740 kb and 21 loss CNVs ranging in size from ~309-1532 kb that overlap CNVs implicated in the etiology of 24 known syndromes and genomic disorders (Additional Table S9). Fourteen individuals had CNVs, including 1 loss (~442 kb) and 13 gains (~414-537 kb), that overlap with the genomic region implicated in Prader-Willi /Angelman syndromes (Type 1), which is caused by a ~5.69 mb deletion on chromosome 15. Thirty-two individuals had CNVs, including 7 losses and 25 gains, ranging in size from ~300-485 kb that overlapped with the region implicated in ATR-16 syndrome, which is caused by a 775 kb deletion on chromosome 16.

## Discussion

The vast majority of existing genetic variation analyses have been performed on individuals of European descent (20). These types of analyses have resulted in an incomplete view of the genetic variation across populations and hindered the understanding and discovery of associations between diseases and genetic variations in non-European populations. To better catalog the full extent of genetic variation across human populations, targeted analyses of genetic variation in under-represented populations are needed. Several recent studies have undertaken such analyses, including of single-nucleotide variations (SNVs), small insertion-deletions (indels), and copy number variations (CNVs) in under-represented populations including people of African, Asian, Latinx and Native American ancestry (21,28–36). Here, we present a catalog of genome-wide copy number variations in a large cohort of healthy individuals of African ancestry.

One of the earliest studies reporting CNVs in a population of African descent was an analysis of 385 individuals of African-American ancestry, which identified 1362 total CNVs (28). Compared to the results we show here, this study used a lower resolution array platform that contained fewer probes, which resulted in a relatively small number of CNVs being identified (28). Over the years, additional studies of individuals from diverse populations, including of African descent as part of 1000 Genomes Project, reported an increasing number of CNVs (>50000) (37–40). Most recently, CNVs and other structural variants (> 400000) in 4937 individuals of African and African-American ancestry were reported as part of the Genome Aggregation Database (gnomAD) (21,24), and novel CNVRs were identified by Nyangiri and colleagues (25).

In this study, we detected > 400000 CNVs including 355027 losses and 61850 gains distributed across all autosomes. The ability to detect CNVs varies between platforms, as SNP-based array platforms are more likely to under-estimate gain CNVs than are array CGH platforms (41,42). Therefore, the number of detected losses is usually higher than the number of detected gains. The vast majority of our CNVs overlap with CNVs in existing databases, including the Database of Genomic Variants (DGV) and gnomAD, indicating that the CNVs we detected are likely true positives. CNVRs observed in our dataset, but not in other existing databases are likely to be either specific to Africans or rare in other populations, underscoring the importance of genetic reference datasets derived from diverse ancestral populations.

We observed a considerable overlap between genes within common CNV blocks and genes previously implicated in developmental disorders curated within the DDG2P Database. These observations raise the possibility that dosage alteration of these genesis either not pathogenic or incompletely penetrant in people of African ancestry. Additionally, of the 866 large CNVs (≥ 300 kb) we identified, 87 overlap with CNVs previously implicated in syndromes catalogued in DECIPHER (27). Thirty of these (34%) are in the same direction (loss or gain) as observed in these known syndromes but are smaller than the pathologic CNVs. One potential explanation for this could be that the region responsible for the clinical outcomes observed in syndromic patients is smaller and our data may allow further refinement of the critical region for these syndromes. Alternatively, these results may also point to variable expressivity and/or reduced penetrance of CNVs in these regions in Africans. These findings underscore the need for population specific CNV datasets for comparison in order to determine the impact of CNVs on clinical outcomes observed in patients (43,44).

A recent study (45) showed that the African “pan-genome”, built using sequence data from 910 individuals of African descent, contained ~10% more DNA not present in hg38 (GRCh38), suggesting that the current reference genome may not fully represent genomic variation in diverse human populations. This suggests the need for *de novo* sequencing of a large number of genomes from African and other under-represented populations, in order to comprehensively assess genomic variation within and between diverse populations.

## Conclusion

The increasing number of African samples being analyzed as part of the 1000 Genomes Project, gnomAD, and several other projects continues to improve our understanding of genetic diversity in this population. However, it is clear that the level of genomic diversity that exists within African subpopulations will require additional, larger datasets in order to capture all the existing genomic variation (46–50). Our data contributes to this effort by providing a rich dataset of CNVs observed in a large cohort of Bantu Africans.

## Materials and Methods

### Sample Description – Populations

Our study cohort included 3631 Bantu African children aged 3-21 living in Mwanza, Tanzania, a region with a population that is both genetically and environmentally relatively homogeneous (51). The subjects were previously genotyped at the Center for Inherited Disease Research (CIDR) as part of the NIDCR FaceBase1 initiative. Age, sex, height, weight, school, and detailed parental and grandparental ethnicity and tribe were collected for all subjects. Individuals with a birth defect or having a relative with orofacial cleft were excluded (51). Genotyping using the Illumina HumanOmni2.5Exome-8v1_A (also referred to as Infinium Omni2.5-8) beadchip array and quality control (QC) was described previously (51,52).

### CNV Detection and Analysis

Signal intensity data (*.idat) files were processed and normalized using Illumina GenomeStudio software. The FinalReport files were used as the raw data to perform CNV calling with three CNV calling algorithms: PennCNV (version 1.0.1) (53), DNAcopy (version 1.46.0), (54) and VanillaICE (version 1.32.2), (55). Both PennCNV and VanillaICE implement Hidden Markov Models (HMM), whereas DNAcopy implements a Circular Binary Segmentation (CBS) algorithm. GC correction was performed for PennCNV using the built-in function, and the R/Bioconductor package ArrayTV (version 1.8.0) (56) was used to perform GC correction for DNAcopy and VanillaICE. Codes used to run the algorithms are available at GitHub (57). Individuals with a total number of CNVs ≥ 3 standard deviations above the cohort mean were removed from further analysis based on previously established criteria (58). In all, 168 individuals were excluded from further analysis: 70 duplicate samples, 97 individuals with a total number of CNVs ≥ 3 standard deviation of the cohort mean, and one individual who had 0 CNVs after applying analysis pipeline thresholds described in Fig. 1. All subsequent analyses were performed on the remaining 3463 individuals and all CNV coordinates are based on NCBI build37/hg19.

CNV calls with fewer than five probes and < 1000 bases in size were removed, followed by those with DNAcopy log-ratio between −0.1 and 0.1 (a threshold determined by a plateau plot in the DNAcopy R package that shows the copy number across the genome), and PennCNV calls with confidence score < 10 (recommended threshold by the developers of PennCNV) (Fig. 1). We used the *intersect* function in BEDTools v2.25 (59) to determine the proportion of overlap between CNV coordinates and genomic elements. CNV calls from two or more algorithms that overlap by 50% or more were considered concordant and included for further analyses. Next, CNV calls overlapping the centromere, telomere, or ≥ 50% with segmental duplications were removed.

PennCNV calls with copy numbers of 0 and 1 were annotated as copy number loss, 2 as diploid copy number, and 3, 4, 5 and 6 as copy number gain; VanillaICE calls with copy numbers of 1 and 2 were annotated as copy number loss, 3 and 4 as diploid copy number, and 5 and 6 as copy number gain; DNAcopy segments with log-ratio ≥ 0.1 were annotated as copy number gain, and log-ratio ≤ −0.1 as copy number loss.

CNV calling with PennCNV from genotype data using high-density SNP arrays often results in the artificial splitting of larger CNVs (i.e. > 500 kb) into multiple smaller CNVs (53). Therefore, we merged adjacent CNVs of the same type (i.e., loss or gain) in the same individual using an approach described previously (53). Briefly, for three adjacent genomic regions A, B, and C, where A and C represent two CNVs of the same type separated by a region B, the length of B was divided by the total length of all three segments (A+B+C). If this fraction was ≤ 15 %, then three regions were merged into one CNV. This approach was used to generate a list of CNVs that passed quality metrics and filtering criteria in individual samples from the Bantu cohort (Additional Table S1).

### In Silico Quality Assessment of CNVs

To assess the quality of CNV calls in the Bantu population, we compared the overlap of CNVs in the Bantu population with the Database of Genomic Variants (DGV) Gold Standard (GS) variants (23). DGV GS variants are a curated set of variants from a select number of studies with high resolution and high quality, which were evaluated for accuracy and sensitivity. Therefore, an overlap with DGV GS variants indicates that our CNV calls are likely true positives. To assess whether the overlap was more than expected by chance, we permuted the genomic locations (n = 1000) using the *shuffle* function in BEDTools v2.25 (59). Permutation tests were performed within each chromosome with the same number and size distribution of CNVs observed in the Bantu population as recommended for genomic elements that are unevenly distributed across the genome (60).

### CNV Regions (CNVRs)

CNV regions (CNVRs) were generated by merging all overlapping CNVs of the same type (i.e. loss or gain) from multiple individuals in our cohort, using the *merge* function in BEDTools v2.25 (59). This resulted in a list of loss-only and gain-only CNVRs, which were further merged into overlapping CNVRs of all types (Additional Table S2).

### Comparison to other CNV Datasets

We compared Bantu CNVRs to variants obtained from DGV (release date 2020-02-15) (23), the Genome Aggregation Database (gnomAD v2.1) (21,24), African CNVR (25) and CNVs identified in low-mappability regions (26). DGV CNVs dataset were downloaded from DGV website (61). gnomAD SV 2.1 sites BED file was downloaded from Broad Institute website (62), which were filtered by SV Type and SV Filter, and only “DEL”, “DUP”, “CN” SV types, and SVs with “PASS” SV Filter were included. The CNV dataset for low-mappability regions obtained from Monlong and colleagues’ publication additional material section (26). CNVs obtained from tumor samples were excluded. CNVRs were generated using a similar approach as described above, and we then compared to the list of Bantu CNVRs to identify overlap.

### CNV Blocks

We generated a list of ‘CNV blocks’ to obtain a more accurate count of the number of times any given CNV was observed in a set of unrelated individuals in our cohort (the description of unrelated individuals is explained in Ref. 51). First, all overlapping CNVs localizing to a given genomic region were aligned as shown (Fig. 4-a,b). The largest region encompassed by these overlapping CNVs (A-D in Fig. 4) was segmented by start and end coordinates of individual CNV calls (A-K in Fig. 4), which resulted into multiple CNV blocks (A-E, E-C, C-J in Fig. 4, Additional Table S5). An example for CNV blocks is represented in Fig. 4b. We then counted the number of times each CNV block was observed in unrelated individuals in our cohort. Based on these counts, CNV blocks were categorized into four groups: CNV blocks observed in ≥ 5% (common CNV blocks), ≥ 1 and < 5% (low frequency CNV blocks), ≥ 0.1 and < 1% (rare CNV blocks), and ≤ 0.1 % (very rare CNV blocks).

### CNVs in Regions Associated with Disease

To assess which CNVs from our cohort overlap genes associated with developmental disorders, we identified overlap (at least 1 bp) of our common (≥ 5%), low frequency (> 1 - < 5%), and rare (> 0.1 - < 1%) CNV blocks with genes catalogued in the Developmental Disorders Genotype-Phenotype Database (Additional Table S4) (DDG2P, 63), compiled based on known implication in disease etiology. The following “STATUS” categories were included in the analysis: Confirmed developmental disorder (DD) Gene, Probable DD Gene, Possible DD Gene, and Both DD and IF (incidental finding). We determined the degree of overlap between using a bi-directional approach; first we calculated how much of the CNV block overlapped with gene (CNVvsGeneOverlap% in Additional Table S6) and then how much of the gene overlapped with the CNV block (GenevsCNVOverlap% in Additional Table S6).

To assess whether large CNVs from our cohort overlap loci associated with genomic disorders, we first generated a list of 866 large CNVs (≥ 300 kb) observed in our cohort (Additional Table S7). We then determined the proportion overlap of these CNVs with known CNVs previously implicated in the etiology of syndromes and genomic disorders catalogued in The DatabasE of genomiC variation and Phenotype in Humans using Ensembl Resources (27,64) (Additional Table S8). DECIPHER is an expert-curated database of microdeletion and microduplication syndromes in developmental disorders.

## List of Abbreviations

CNV: Copy number variation
CNVR: Copy number variant region
SNV: Single nucleotide variation
DGV: Database of Genomic Variants
GS: Gold Standard
DDG2P: Developmental Disorders Genotype-Phenotype Database
gnomAD: Genome Aggregation Database
SE: Standard Error
IQR: Interquartile

## Declarations

### Ethics Approval and Consent to Participate

Written informed consent was obtained for all study subjects or their parents, as appropriate. This study was approved by the Colorado Multiple Institutional Review Board, the University of Calgary Institutional Review Board, and the National Institute for Medical Research (Tanzania).

### Data Availability

CNV data have been deposited with FaceBase (https://doi.org/10.25550/1-7330).

### Conflict of Interest

None

### Funding

This work was supported in part by grant # DE025363 from the National Institutes of Health to T.H.S.

### Author Contributions

T.H.S and A.E.H conceived the study. F.Y., D.A. and H-C. Y. performed the copy number analysis and data interpretation. R.A.S led the original study that collected samples and conducted the genotyping study of the Bantu cohort. J.C, S.A.S and R.A.S provided the raw intensity data and other relevant information on the sample used in this study. F.Y., M.N., A.E.H. and T.H.S. drafted the manuscript which was read and critically revised by all authors. Final approval of the version to be published was given by F.Y., M.N., D.A., H-C.Y., J. C., S. A. A., R. A. S., A. E. H., and T.H.S.

## Acknowledgements

We would like to thank the FaceBase Consortium for providing the genotyping data used in this study. University of Colorado Anschutz Medical Campus Department of Biochemistry and Molecular Genetics’ research cluster was used to perform analyes. The genotype data used for CNV detection were previously deposited in the Database of Genotypes and Phenotypes (dbGaP: http://www.ncbi.nlm.nih.gov/gap; dbGaP study accession: phs000622.v1.p1). This study makes use of data generated by the DECIPHER community. A full list of centres who contributed to the generation of the data is available from http://decipher.sanger.ac.uk and via email from. Funding for the project was provided by the Wellcome Trust.

---

**Additional File 1:** AdditionalFiles_TableS1.xlsx. Title: The list of CNVs. Description:

The list of CNVs detected in our study.

**Additional File 2:** AdditionalFiles _TableS2.xlsx. Title: The list of CNVRs.

Description: The list of CNVRs identified in our study.

**Additional File 3:** AdditionalFiles_TableS3.xlsx. Title: Novel CNVs observed in our cohort. Description: The list of novel CNVs detected in our study.

**Additional File 4:** AdditionalFiles_TableS4.xlsx. Title: The list of DDG2P genes used in our analysis. Description: The list of DDG2P genes used in our analysis to detect genes overlapping with Bantu CNV blocks.

**Additional File 5:** AdditionalFiles_TableS5.xlsx. Title: The list of CNV blocks. Description: The list of Bantu CNV blocks identified in our study.

**Additional File 6:** AdditionalFiles_TableS6.xlsx. Title: CNV blocks overlapped with DDG2P genes. Description: The list of Bantu CNV blocks overlapped with DDG2P genes.

**Additional File 7:** AdditionalFiles_TableS7.xlsx. Title: The list of large (>300kb) CNVs observed in unrelated individuals in our cohort. Description: The list of large CNVs identified in our study.

**Additional File 8:** AdditionalFiles_TableS8.xlsx. Title: The list of CNVs associated with DECIPHER Syndromes used in our analysis: Description: The list of DECIPHER CNV syndromes used in our study.

**Additional File 9:** AdditionalFiles_TableS9.xlsx. Title: CNVs associated with

DECIPHER syndromes overlapping large CNVs observed in our cohort. Description: The list of DECIPHER CNV syndromes overlapped with DECIPHER CNV syndromes.

## Notes

### Competing Interest Statement

The authors have declared no competing interest.

